# β-arrestin recruitment facilitates a direct association with G proteins

**DOI:** 10.1101/2025.06.24.661366

**Authors:** Claudia Y. Lee, Jeffrey S. Smith, Taylor Kohlmann, Emily M. Meara, Uyen Pham, Frank Kwarcinski, Andrew N. Dates, Issac Choi, Ari S. Hilibrand, Abigail Gillikin, Stephen C. Blacklow, Gregory G. Tall, Andrew C. Kruse, Sudarshan Rajagopal

## Abstract

G protein-coupled receptors (GPCRs) are targets for almost a third of all FDA-approved drugs. GPCRs are known to signal through both heterotrimeric G proteins and β-arrestins. Traditionally these pathways were viewed as largely separable, with G proteins primarily initiating downstream signaling while β-arrestins modulate receptor trafficking and desensitization in addition to regulating their own signaling events. Recent studies suggest an integrated role of G proteins and β-arrestins in GPCR signaling, however the cellular and biochemical requirements for G protein: β-arrestin interactions remain unclear. Here we show that G proteins and β-arrestins can directly interact. Through utilization of β-arrestin-biased receptors and artificially enforced β-arrestin relocalization, we demonstrate that recruitment of β-arrestin to the plasma membrane is sufficient to interact with the G protein Gαi. Using purified proteins, we show that Gαi directly interacts with β-arrestin. In addition, we find that Gαi family members differ in their degree of association with β-arrestin, and that a large degree of this selectivity resides within the alpha helical domain of Gαi. These findings delineate the cellular and biochemical conditions that drive direct interactions between G proteins and β-arrestins and illuminate the molecular basis for how they work together to effect GPCR signaling.

## Introduction

GPCRs are critical regulators of human physiology, allowing cells to detect and respond to a wide range of stimuli ranging from peptides and small molecules to odorants and photons of light. Canonically, GPCRs are known to initiate signaling through two primary signaling pathways (1) heterotrimeric G proteins and (2) β-arrestins^1–3^. In G protein signaling, a conformational change in a GPCR allows it to catalyze exchange of Gα-bound GDP for GTP. The activated Gα then dissociates from the receptor and from the Gβγ subcomplex. Gα and Gβγ proceed to regulate the activity of downstream signaling proteins such as adenylyl cyclase, inositol trisphosphate receptor, and protein kinase C, as well as modulate ion channel activity^1^. In the canonical β-arrestin pathway, GPCR activation results in phosphorylation of serine and threonine residues, primarily by GPCR Kinases (GRKs) on the flexible C-terminal tail and larger intracellular loops. Phosphorylation of intracellular residues promotes β-arrestin binding to the receptor core, which sterically prevents Gα binding to the receptor, leading to desensitization of G protein signaling^4,5^. This interaction also facilitates plasma membrane anchoring of β-arrestins, despite the β-arrestins lacking lipid membrane-anchoring post translational modifications^6^. In addition, β-arrestins facilitate receptor internalization and can directly scaffold several other proteins, such as Src, extracellular-signal regulated signal kinase (ERK), and Raf, to initiate β-arrestin-dependent signaling events^7–13^. Ligands have been identified that preferentially activate either the G protein or the β-arrestin signaling pathways. It was thought that such ‘biased-agonists’ could reduce certain ontarget, but unintended, adverse effects while enhancing therapeutic efficacy. For example, at the µ-opioid receptor (MOR), the G protein-biased ligand TRV130 (oliceridine)^14^ is now FDA-approved for the treatment of acute pain. In addition to ligand bias, some receptors display intrinsic signaling preferences, a phenomenon referred to as ‘receptor bias’. For instance, the atypical chemokine receptor ACKR3 preferentially engages β-arrestins over G proteins^15^.

Despite promising initial results, the widespread clinical success of biased signaling strategies has been more limited than expected. In addition, as signaling assay sensitivity and specificity has improved, the extent of cross-talk between signaling pathways regulated by both G proteins and β-arrestins is now appreciated^16,17^. Numerous signaling studies suggest an interrelated role of G proteins and β-arrestins in executing the multifaceted signaling signature of each GPCR. For example, concurrent discoveries in 2009 found that prolonged binding of β-arrestin to the parathyroid hormone receptor type 1 or activation of thyroid stimulating hormone receptor in endosomes prolongs cAMP production, implicating β-arrestin-regulated signaling in promoting endosomal G protein activity^18,19^. Further evidence for coordinated G protein and β-arrestin signaling has come from structural studies using negative stain and cryo-electron microscopy, which revealed a signaling “megaplex” involving a β2-Adrenergic Receptor/Vasopressin 2 Receptor chimera simultaneously engaging Gαs at the receptor core and β-arrestin at the phosphorylated C-terminal tail^20,21^. Gαs can also direct GRK isoform recruitment, thereby modulating β-arrestin recruitment and alter β-arrestin conformation in cells as assessed by β-arrestin biosensors^22^. Consistent with cross talk between β-arrestin and G protein signaling, studies have recently demonstrated that G proteins and β-arrestins can associate following activation of multiple GPCRs^23^. These complexes differed from previously reported G protein:β-arrestin cross talk as 1) this association was specific for inhibitory G protein (Gαi), even among receptors which do not classically signal through Gαi, and 2) these complexes formed primarily at the plasma membrane. Gαi:β-arrestin complexes can coordinate with secondary effectors, such as ERK and clathrin adapter AP-2^23,24^.

In order to better understand and untangle the complex interplay of G protein and β-arrestin signaling, we sought to define the biochemical and cellular requirements for G protein:β-arrestin interactions. In cells, we found that recruitment of β-arrestin to the plasma membrane, either via β-arrestin-biased receptors or chemically inducible heterodimerization, was both necessary and sufficient to promote Gαi:β-arrestin complex formation. In contrast, activation of Gαi alone, using G protein-biased agonists or aluminum fluoride, did not induce complex formation. We confirm this by leveraging the FKBP/FRB heterodimerization system as an artificial, receptor-independent method to translocate β-arrestin to the plasma membrane. Using purified proteins, we show that Gαi can directly interact with β-arrestin, and that this interaction is not dependent on the nucleotide state of the Gα subunit. We further demonstrate that β-arrestin binding does not modulate nucleotide exchange by Gαi proteins. Finally, we find that the extent of β-arrestin association varies between different Gαi isoforms and reveal that regions within the alpha helical domain plays a crucial role in β-arrestin binding in living cells. Together, our experiments reveal the biochemical and cellular conditions that promote direct interactions between G proteins and β-arrestins.

## Results

### The M3 muscarinic receptor facilitates Gαi:β-arrestin complex formation

We first re-established our previous findings of Gαi:β-arrestin and tested if agonist treatment of the M3 muscarinic receptor (M3R) could lead to G protein:β-arrestin complex formation in HEK293T cells by NanoBit complementation. M3R primarily signals through the G protein Gαq, but can also signal through Gαi and Gαs^25–27^. Agonist treatment of the M3R facilitated observable and robust associations of β-arrestin with Gαi, but not Gαq nor Gαs in cells (Figure 1A,B). Multiple muscarinic agonists facilitated Gαi:β-arrestin complex formation (Figure 1C). Pretreatment with the muscarinic antagonist tiotropium prevented complex formation and increasing concentrations of agonist increased Gαi:β-arrestin association, indicating a receptor-specific effect (Figure 1D,E).

**Figure 1.**
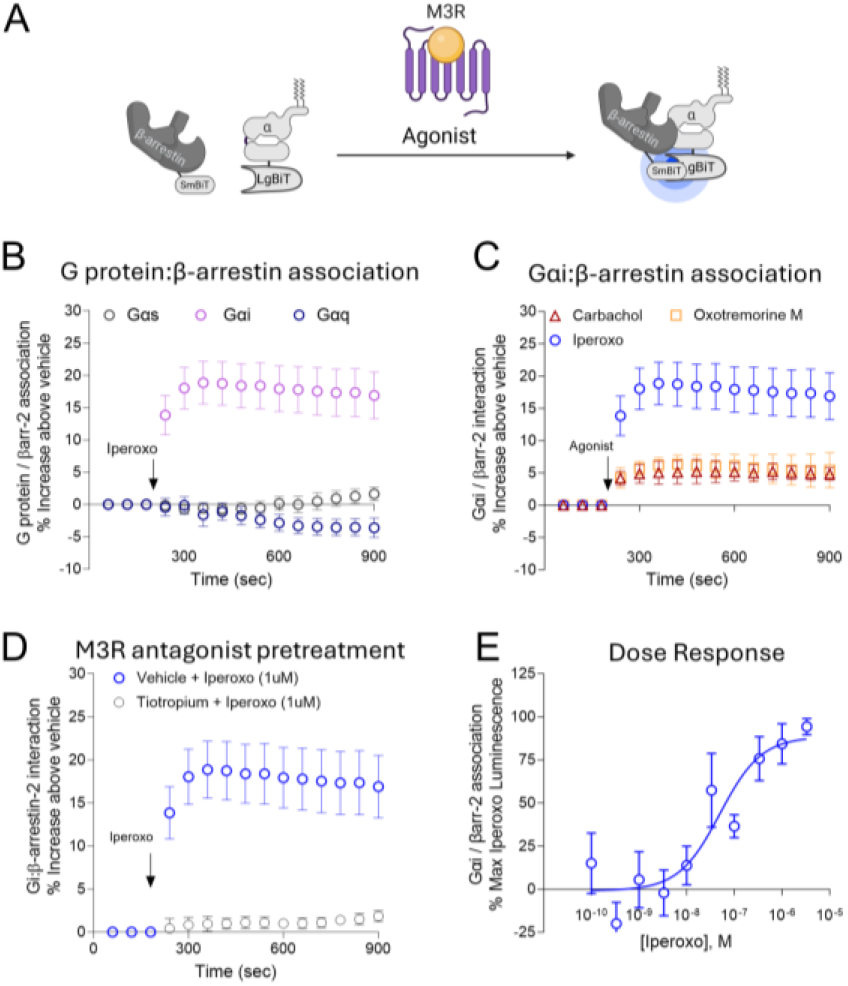
M3R activation promotes Gαi:β-arrestin association. **(A)** Cartoon of split luciferase assay. **(B)** HEK 293T cells transiently transfected with the M3R, the indicated Gα-LgBiT, and SmBiT-β-arrestin-2 were treated with the muscarinic agonist iperoxo (1 µM). **(C)** Cells transiently transfected with the M3R, Gαi1-LgBiT, and SmBiT-β-arrestin-2 were treated with the indicated muscarinic agonist iperoxo (1 µM), carbachol (100 µM), or oxotremorine-M (10 µM). **(D)** Cells were pretreated with either tiotropium (10 µM) or vehicle for 10 minutes and subsequently treated with the indicated iperoxo (1 µM). **(E)** Cells transiently transfected with the M3R, Gαi1-LgBiT, and SmBiT-β-arrestin-2 were treated with iperoxo at the indicated concentration. N=3 replicates per condition. Data shown are mean ± SEM.

### β-arrestin-biased GPCRs promote Gαi:β-arrestin complex formation

The current paradigm for GPCR activation generally posits that the receptor active state confers conformational changes in both G proteins and β-arrestin to initiate their signaling. However, it is also known that G proteins and β-arrestins may have receptor-independent mechanisms of activation^28–30^. To assess whether Gαi activation was required for Gαi:β-arrestin association, we utilized three β-arrestin-biased receptors that are largely incapable of activating G proteins, including Atypical Chemokine Receptor 3 (ACKR3), Angiotensin II type I Receptor AAY mutant (AT1R-AAY), and the Dopamine 2 Receptor ARB mutant (D2R-ARB)^15,31,32^. AT1R-AAY and D2R-ARB contain mutations in their DRY motifs that disrupt G protein activation, causing them to signal primarily via β-arrestin-mediated pathways, while ACKR3 is a naturally occurring β-arrestin-biased receptor^15,33^. For AT1R-AAY, we tested both endogenous ligand angiotensin II (AngII) and known β-arrestin-biased agonist, TRV120023 (TRV023)^34^. For ACKR3, we tested multiple ligands, including two small molecules that selectively target ACKR3^35,36^. Despite the lack of G protein activation at these receptors, we observed that each β-arrestin-biased receptor promoted Gαi:β-arrestin complex formation (Figure 2A-D). Furthermore, we showed at ACKR3 these complexes form in the absence of G protein recruitment to the receptor (Figure 2B, Supp. Figure 1C-D). These results demonstrate that Gαi:β-arrestin complex formation can occur independently of classical receptor-mediated G protein activation.

**Figure 2.**
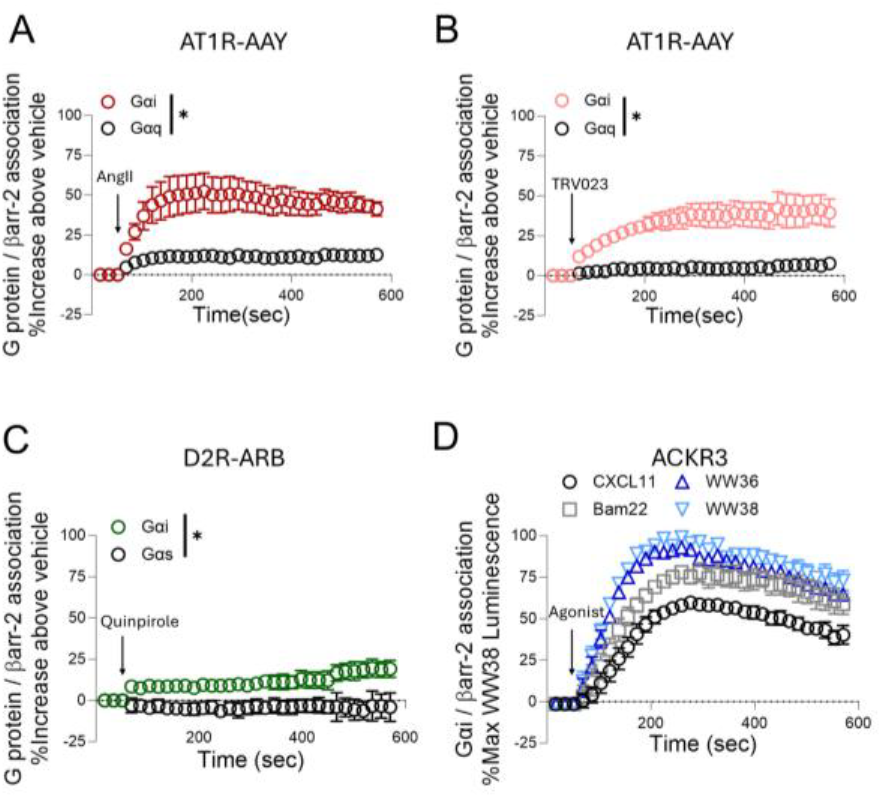
β-arrestin-biased GPCRs promote Gαi:β-arrestin association. **(A)** HEK 293T cells transiently transfected with the indicated β-arrestin-biased receptor, the indicated Gα-LgBiT, and SmBiT-β-arrestin-2 (A) AT1R-AAY stimulated with 1 µM AngII **(B)** AT1R-AAY stimulated with 10 µM TRV023. **(C)** D2R-ARB stimulated with 1 µM Quinpirole. **(D)** ACKR3 stimulated with 100nM of its endogenous ligand, CXCL11 (C-X-C Ligand 11), 1 µM Bam22, 1 µM WW36, and 1 µM WW38. For panel A-B *P<0.0001 by two-tailed t test comparing area under curve (AUC) for Gαi-AngII versus Gαq-AngII (A) and Gαi-TRV023 versus Gαq-TRV023 (B). For panel C *P<0.005 Gαi versus Gαs by two-tailed t test comparing AUC. n=3 for all conditions and represents independent plate replicates. Graphs depict the percent increase over vehicle treatment normalized to max Gαi signal (A-C) or percent ACKR3 max WW38 signal (D). Data shown are mean ± SEM.

### G protein-biased ligands fail to promote Gαi:β-arrestin complex formation

To complement experiments with β-arrestin-biased receptors, we then tested whether activating the traditional G protein pathway without β-arrestin recruitment was sufficient to facilitate Gαi:β-arrestin association. The Gαi-coupled MOR has several well characterized agonists available including balanced agonists such as [D-Ala2, N-MePhe4, Gly-ol]-enkephalin (DAMGO) and fentanyl, as well as G protein-biased agonists including oliceridine (also known as TRV130) and PZM2137. We first validated that oliceridine and PZM21 activated G proteins and did not promote observable β-arrestin recruitment, as expected, while DAMGO and fentanyl promoted both G protein activation and β-arrestin recruitment (Figure 3A-E). We proceeded to test these G protein-biased agonists for their ability to facilitate Gαi:β-arrestin complex formation. In contrast to the balanced ligands which activated both G protein and β-arrestin pathways (Figure 3F-H), oliceridine and PZM21 failed to promote Gαi:β-arrestin complex formation. These findings suggest that G protein activation alone is not sufficient for Gαi:β-arrestin complex formation and highlight the requirement of β-arrestin recruitment in mediating this interaction.

**Figure 3.**
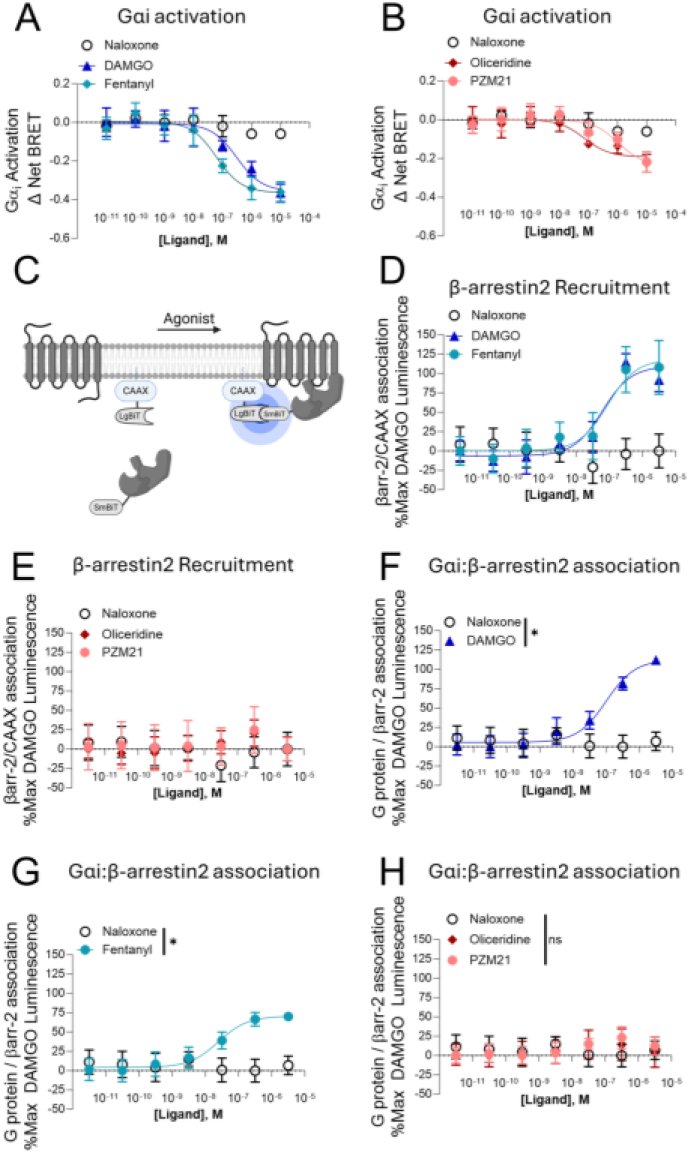
G protein-biased ligands do not promote Gαi:β-arrestin complex formation. HEK293T cells overexpressing the Gαi TRUPATH BRET sensor with **(A)** balanced ligands DAMGO, and Fentanyl, (n=3) or **(B)** G protein biased ligands Oliceridine and PZM21 (n=3). The antagonist Naloxone was used as a negative control. **(C)** Illustration of by-stander NanoBiT recruitment assay used to monitor β-arrestin-2 recruitment upon MOR activation. Briefly, LgBiT is localized to the plasma membrane by a CAAX tag. Upon activation of MOR, the β-arrestin-2 translocates to the plasma membrane to produce luminescence. **(D)** Balanced ligands, DAMGO (n=6) and Fentanyl (n=3), promote β-arres-tin-2 recruitment to MOR. **(E)** G biased ligands, Oliceridine (n=5) and PZM21(n=5), do not recruit β-arrestin-2 to MOR. Naloxone was used as a negative control (n=6) in D-E. **(F-H)** Split luciferase Gαi-LgBiT:smBiT-β-arrestin-2 association upon stimulation with DAMGO (n=9) **(F)**, Fentanyl (n=7) **(G)** but not G protein biased ligands Oliceridine and PZM21 (n=4) **(H)**. Naloxone (n=7) was used as a negative control (F-H). For panel F,G, and H, a two-way ANOVA with Bonferroni post hoc test was performed to compare the effect of DAMGO, Fentanyl, Oliceridine, or PZM21 with the negative control, Naloxone. *P<0.05 (F-G), ns P>0.05 (H). All graphs denote the mean signal ± SEM and are normalized to max DAMGO signal.

### β-arrestin recruitment to the plasma membrane is sufficient to form Gαi:β-arrestin complexes

Since β-arrestin-biased receptors, but not G protein-biased ligands, were capable of promoting Gαi:β-arrestin complex formation, we next tested if β-arrestin recruitment to the plasma membrane in the absence of GPCR agonist treatment was sufficient to form Gαi:β-arrestin complexes. To do this, we utilized a heterodimerization system comprised of FK506 Binding Protein (FKBP) and FKBP Rapamycin Binding domain (FRB) to force β-arrestin to the plasma membrane^38^. In this system we added a plasma membrane-localization Lyn11 tag to FRB and placed a FKBP onto the C-terminus of β-arrestin. In doing so we could translocate β-arrestin to the plasma membrane by the addition of rapamycin and independently of GPCR activation (Figure 4A). Transiently transfecting these constructs in HEK293T cells followed by rapamycin treatment led to the formation of Gαi:β-arrestin, but not Gαs:β-arrestin complexes (Figure 4B-C), suggesting that plasma membranelocalized β-arrestin is sufficient to promote selective Gαi interaction even in the absence of GPCR agonism.

**Figure 4.**
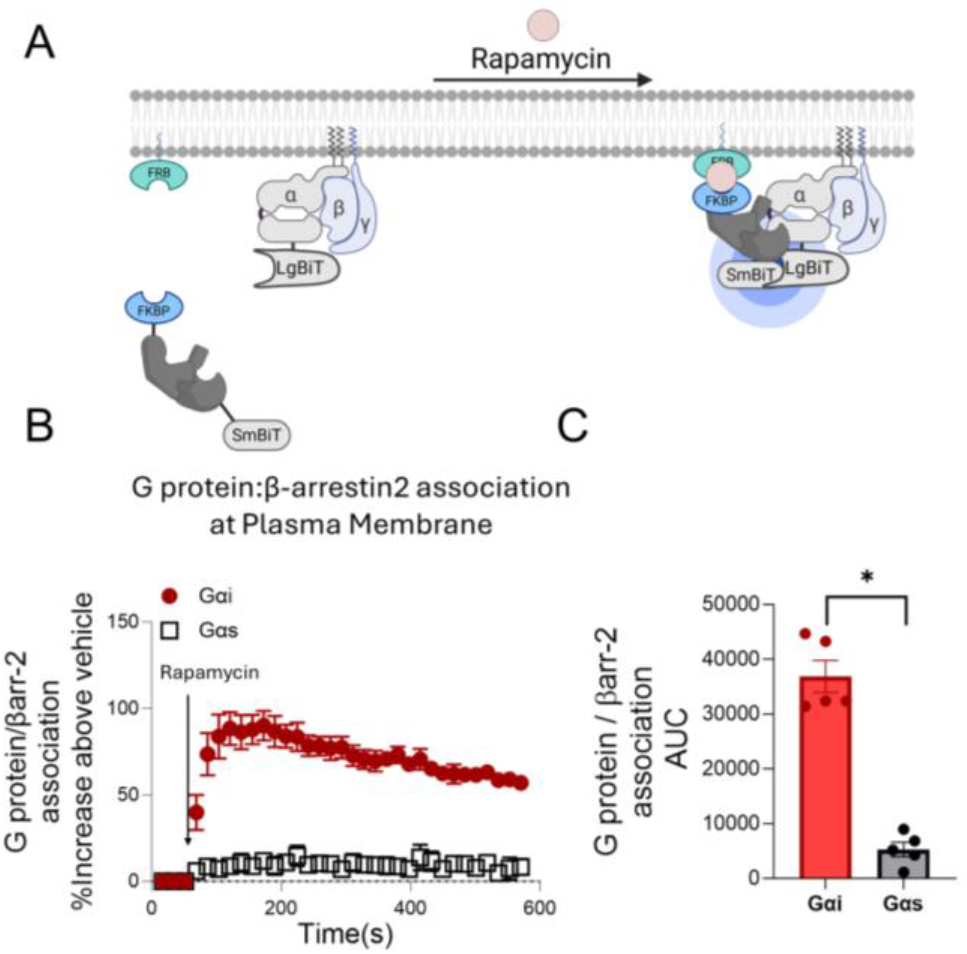
Translocation of β-arrestin to plasma membrane is sufficient to promote Gαi:β-arrestin association. **(A)** Schematic of NanoBiT assay monitoring Gαi:β-arrestin association upon induced translocation β-arrestin-2 to the plasma membrane. Upon addition rapamycin (500 nM), FKBP and FRB heterodimerize, forcing translocation of β-arrestin to the plasma membrane independent of receptor activation. Gαi:β-arrestin complexes are assessed via production of luminescence from reconstituted luciferases. **(B)** Gαi:β-arrestin complex formation following addition of rapamycin. **(C)** AUC analysis of (B). *P < 0.05 by two-tailed t-test with Gαi vs Gαs. n=5 independent replicates for both graphs. Data shown are mean ± SEM.

### β-arrestin directly interacts with Gαi

We then tested if purified β-arrestin and Gαi directly interact. We utilized a C-terminally truncated form of β-arrestin-1 shown to have greater spontaneous receptor-binding activity (β-arrestin-1-393X) and a non-lipidated form of Gαi1 (Supp. Figure 2). Pulldowns of the N-terminal Protein-C tag of β-arrestin led to the coelution of Gαi1, which was not markedly changed when treated with apyrase to remove nucleotides or incubated with excess GDP or GTPγS (Figure 5A). These findings were in line with our previous G protein-biased signaling data and consistent with NanoBRET studies where activation of G proteins by aluminum tetra-fluoride (AlF _4_^-^)^39^ or over expression of a guanine dissociation inhibitor of Gαi, activator of G protein Signaling 3 (AGS3)^40^, did not facilitate Gαi:β-arrestin interactions (Supp. Figure 3). Given this direct interaction, we then tested if purified β-arrestin-1 or β-arrestin-2 could influence the rate of [35S]-GTPγS binding to lipidated or non-lipidated forms of Gαi. Basal Gαi [35S]-GTPγS binding rates were consistent with a previous study^41^. Consistent with the bound nucleotide status of Gαi not overtly influencing the ability to be pulled-down with β-arrestin-1, incubation with β-arrestin-1-393X and β-arrestin-2-392X did not alter the rates of [35S]-GTPγS binding to either myristoylated Gαi2 or unmodified Gαi1, consistent with the pulldowns in which Gαi:β-arrestin complexation was agnostic to nucleotide state of the Gαi protein (Figure 5B).

**Figure 5.**
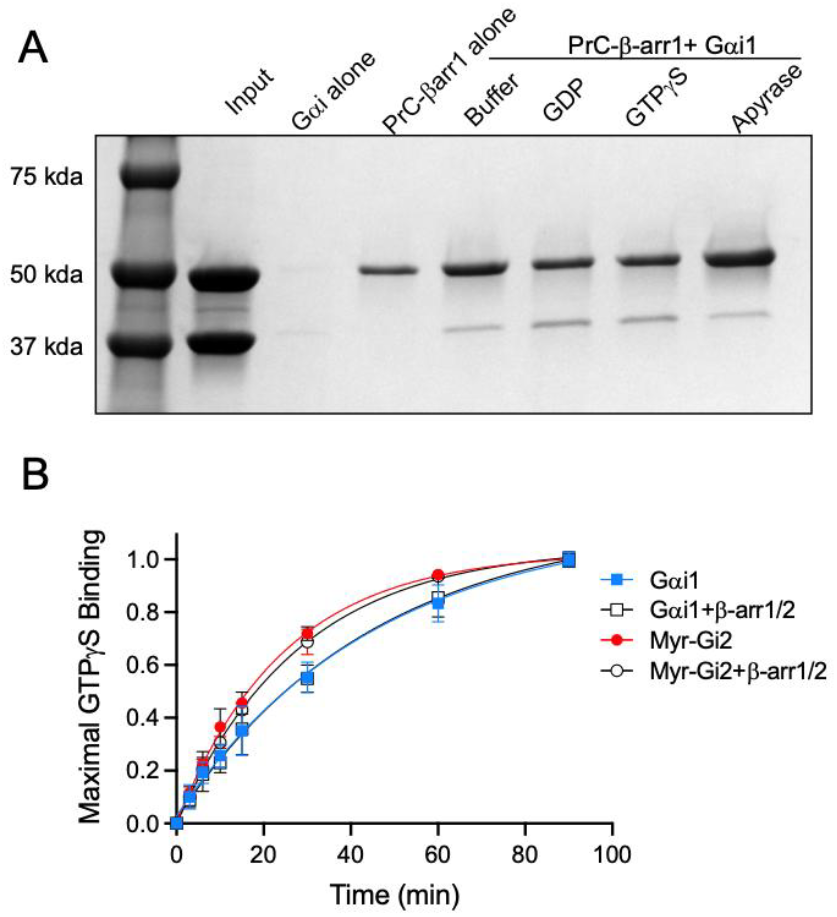
Purified Gαi and β-arrestin directly interact. **(A)** Coomassie gel of a pull down of purified N-terminally protein-c tagged β-arrestin-1-393X with purified non-lipidated Gαi1 incubated either with buffer or various nucleotide conditions (GDP 20 µM, GTPγS 20 µM, or apyrase for the nucleotide-free state). **(B)** [35S]-GTPγS incorporation into non-lipidated Gαi1 or myristoylated Gαi2 in the presence or absence of 4x molar excess of purified β-arrestin-1-393X and β-arrestin-2-393X. For panel A, pull-down representative of three independent experiments. For panel B, N=3 replicates are plotted as maximal GTPγS binding. Error bars are the mean ± S.D.

### Mutagenesis reveals domains of Gαi critical for interaction with β-arrestin

To identify the structural domains within Gαi that are crucial for Gαi:β-arrestin association, we generated chimeras of Gαi and with Gα proteins that do not interact with β-arrestins. We first probed the C-terminal α5 helix, known for its key role in receptor binding and selectivity^42^, by swapping the α5-helix of Gαi into Gαs (Gαs-i-helix 5) and vice versa (Gαi-s-helix 5). Exchange of helix 5 had no effect on β-arrestin binding by either G protein, as evidenced by clear complex formation by Gαi-s-helix5, but not the Gαs-i-helix 5 (Figure 6A), suggesting that the Gαi α5-helix is not the primary location for its interaction β-arrestin.

**Figure 6.**
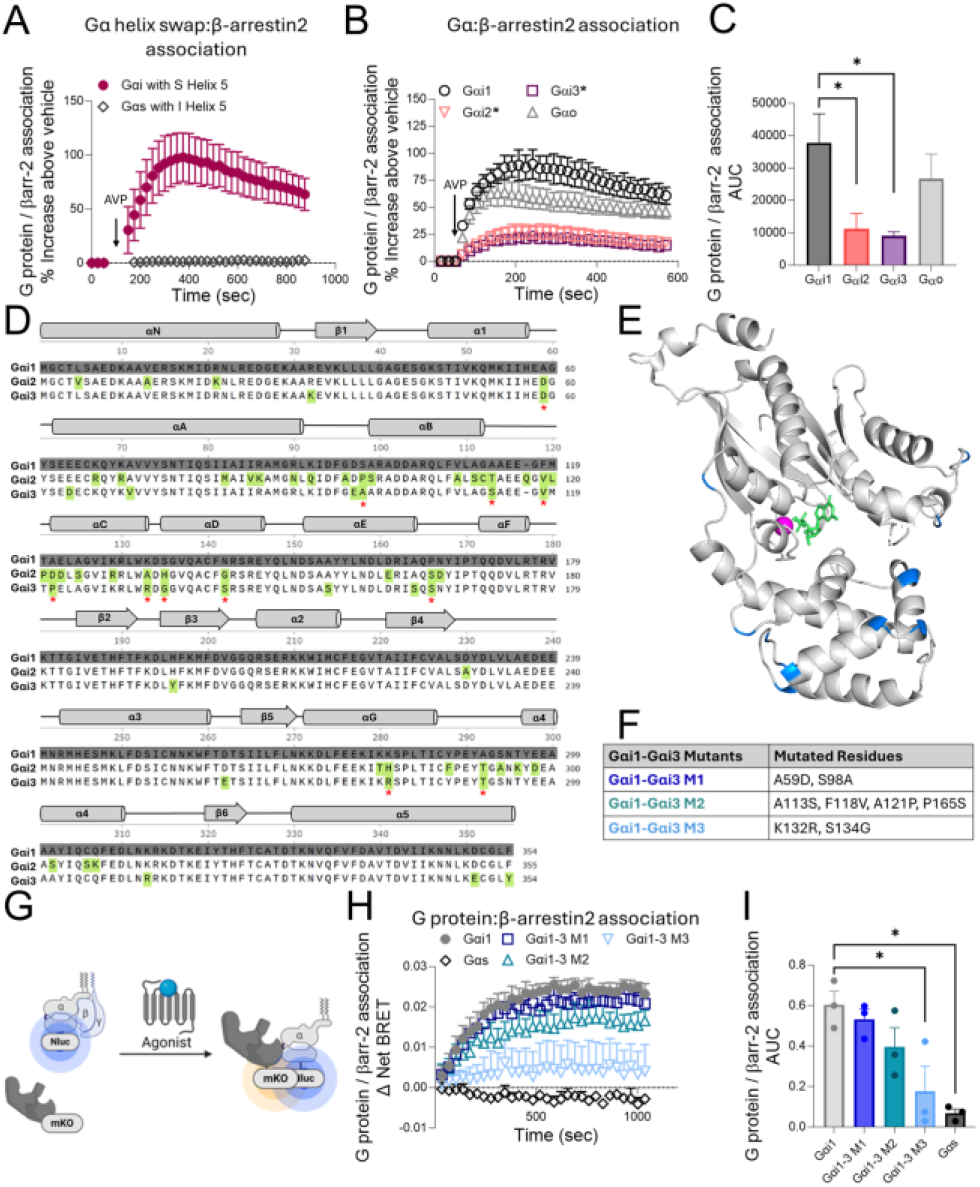
Mutations within Gα helical domain impairs interactions with β-arrestin. **(A)** HEK293T cells were transiently transfected with V2R, SmBiT-β-arrestin-2, and either LgBiT-tagged Gαi with Gαs α5 helix or LgBiT-tagged Gαs with Gαi α5helix and treated with AVP (500 nM) **(B)** LgBiT-tagged Gαo, Gαi1, Gαi2 and Gαi3 were compared with their ability to interact with SmBiT-β-arrestin-2 in cells overexpressing the V2R and treated with AVP (500 nM). (**C)** Area under the curve graph (AUC) made from the kinetic tracing data in panel B. *P <0.05 by one-way ANOVA, Dunnett post hoc test of Gαi1 versus Gαi2, Gαi3, or Gαo. **(D)** Sequence alignment of human Gαi1, Gαi2, Gαi3 with differing residues highlighted with red asterisks. Alignment performed by GproteinDb66,67 **(E)** Structure of inactive Gαi1 (PDB:1BOF^68^) residues distinct between Gαi1 and Gαi2 and Gαi3 are highlighted in green. **(F)** Diagram of the NanoBRET assay for Gαi:β-arrestin complex formation. **(G)** Gαi1 A59D, S98A (M1), Gαi1 A113S, F118V, A121P,P165S (M2), or Gαi1 K132R S134G (M3) were compared to Gαi:β-arrestin by NanoBRET assay. HEK293T cells were transiently transfected with Nluc-tagged Gαi mutants (WT, M1, M2, or M3), β-arrestin-2-mKO, and V2R. Upon activation of V2R with AVP (500 nM), cells were monitored for changes in BRET ratio indicating Gαi:β-arrestin complex formation. **(H)** Area under the curve graph (AUC) made from the kinetic tracing data in panel G and represents Δ Net BRET between AVP and vehicle treated cells. *P<0.05 by one-way ANOVA with Dunnett’s post hoc comparing Gαs versus Gαi1 and Gαi1-3 M3 versus Gαi1. Data in A-B demonstrate % increase luminescence of AVP above vehicle signal. All data shown are from n=3 replicates. Data shown are mean ± SEM.

Another region where Gαi might interact with β-arrestin is the Gα interface with Gβγ. This region is mainly comprised of the Gα Switch 2 region and is well known to bind several proteins including modulators for Gα activity like Ric-8A (resistance to inhibitors of cholinesterase)^43^ and Gα-interacting vesicle-associated protein (also known as Girdin)^30^ as well as downstream effectors like adenylyl cyclase^44^. To test if Gβγ could compete with β-arrestin for binding to Gαi and impact complex formation, we titrated increasing amounts of Gβγ plasmid into HEK293T cells and assayed Gαi:β-arrestin association. Overexpression of Gβγ did not disrupt Gαi:β-arrestin complex formation (Supp. Figure 4A-B). These data suggest that Gβγ and β-arrestin do not compete for a similar Gαi binding site and reiterates our previous observation that Gαi does not require Gβγ for association with β-arrestin (Figure 5A).

To identify other domains within Gαi that could interact with β-arrestin we looked for differences in β-arrestin binding amongst the closely-related members of the Gαi subfamily. Upon activation of V2R, we found Gαi1 and Gαo formed more robust complexes with β-arrestin compared to Gαi2 and Gαi3, which displayed reduced interaction (Figure 6B-C). These differences were not due to differences in basal expression or basal coupling, as lower amounts of transfected Gαi/Gαo still yielded consistent results (Supp. Figure 5). Based on these findings, we hypothesized that residues which differ between Gαi1 and those in Gαi2 and Gαi3 are important to promote Gαi:β-arrestin complexes. To determine which regions potentially contributed to this difference, we aligned the amino acid sequence of Gαi1, Gαi2, and Gαi3. Upon initial inspection, there were a total of 11 residues which differed between Gαi1 and those of Gαi2 and Gαi3. Several of these residues were located within the alpha helical domain of Gαi1 (Figure 6D-E), suggesting that this region of the protein might confer specificity for interactions with β-arrestin. In fact, the alpha helical domain is a known binding site for Regulator of G Protein Signaling (RGS) proteins, which selectively generate distinct RGS-G protein pairs via residue differences across Gαi isoforms in this region^45,46^. To test if these differing residues within Gαi subfamily members conferred the subtype effect for Gαi:β-arrestin, we generated three Gαi mutants in which non-conserved helical domain residues in Gαi1 were substituted with those from Gαi3. These point mutants were grouped by their spatial proximity within Gαi1 structure and linear sequence location and were referred to as Gαi1-Gαi3 M1, Gαi1-Gαi3 M2, and Gαi1-Gαi3 M3 (Figure 6E). All mutants were tagged with a full-length luciferase (NanoLuc) for use in a NanoBRET assay and to ensure expression differences did not account for variation in signal. Upon treating the V2R with agonist, we observed that Gαi1 K132R S134G (Gαi1-Gαi3 M3) within the alpha helical domain greatly decreased the association with β-arrestin (Figure 6F-H). These results highlight the importance of specific residues within the helical domain of Gαi1, such as K132 and S134, for β-arrestin binding. Given that RGS proteins utilize the same residues in their helical domain for G protein binding and confer specificity by the orientational differences of side chains^46^, we further tested whether the entire helical domain is required for interaction with β-arrestin. To do this, we generated a Gαi-Gαs helical domain swap (Gαi-GαsHD) and evaluated Gα:β-arrestin complex formation using the NanoBiT complementation assay. The Gαi-GαsHD swap had diminished ability to interact with β-arrestin (Supp. Figure 6). The totality of mutagenesis experiments suggested that the alpha helical domain of Gαi is a critical structural determinant for β-arrestin interaction.

## Discussion

In this study, we identified the cellular and certain biochemical factors that promote direct interactions between G proteins and β-arrestins. By using β-arrestin-biased receptors and the FKBP/FRB heterodimerization system, we showed that recruitment of β-arrestin to the plasma membrane was sufficient to promote its interaction with G proteins, even in the absence of receptor activation. Activation of G protein alone, either through G protein-biased agonists or aluminum fluoride, was not sufficient to promote significant G protein:β-arrestin association. In cells, we found that this interaction was most robust with Gαi-family proteins relative to other G protein families. With purified proteins, we showed that β-arrestin directly interacts with Gαi and does not require Gβγ. This interaction did not depend on the nucleotide status of the G protein, as Gα in the GDP-bound, GTP-bound, or nucleotide-free states were all capable of interacting with β-arrestin. Consistent with these observations, purified β-arrestins did not alter the rate of GTP binding to Gαi. Sequence alignment of Gαi-family isoforms and subsequent mutagenesis studies localized the likely interaction site with β-arrestin to the alpha helical domain of Gα, a finding supported by other work utilizing hydrogen-deuterium exchange mass spectrometry (submitted). We also found that the C-terminal α5-helix of Gαi, which engages GPCRs, did not confer specificity for Gαi:β-arrestin. This further supports our findings that G proteins interact with β-arrestins in a manner distinct from their binding to GPCRs. While the absence of clear β-arrestin association with either Gαs or Gαq in HEK293 cells is consistent with prior observations^23,24^, we note that the absence of observed association in our assays does not preclude association under other conditions or in other cell types, as other G protein family members have been observed in close proximity with β-arrestin via crosslinking or in the presence of a GPCR^20,47,48^.

Our data supports a working model where plasma membrane-localized β-arrestin can directly interact with the alpha helical domain of Gαi, independent of a GPCR scaffold. β-arrestin did not provide direct guanine exchange factor (GEF) activity as assessed by the measuring the velocity of GTP binding to Gαi. Such an interaction that does not necessarily require a GPCR is consistent with an ‘action-at-a-distance’ signaling model of β-arrestin previously described by Von Zastrow and colleagues^49^, and seemingly consistent with observations of β-arrestin and Gαi in endosomes that lack an associated receptor^50^. Available structural and functional data suggest that β-arrestins adopt a variety of conformational states and binding partners depending on the cellular compartment, function, and lipid environment^51–54^. Indeed, it is now appreciated that endosomal or organellar signaling initiated by G proteins leads to fundamentally distinct cellular responses^55,56^. We speculate that β-arrestins can help facilitate trafficking and localization of G proteins to various intracellular compartments, although further work is necessary to test this hypothesis.

There are limitations of our study which should be considered. Firstly, some of our experiments utilized overexpressed proteins which may alter the basal signaling processes that occur in our cell system. In some cases, such overexpression can result in diminished signaling specificity downstream of GPCRs^57^. Thus, there may be additional cellular components or cofactors which may have been overlooked in our assessment that can promote, or inhibit, G protein and β-arrestin interactions.

Mounting evidence links the spatial and temporal localization of receptors, G proteins, and β-arrestins to divergent signaling events and physiological responses^55,58^. The physiological relevance of such a direct Gαi:β-arrestin interaction is supported by a growing number of reports. For example, the Gαi:β-arrestin interaction is thought to be critical to follicle stimulating hormone function (a receptor that classically couples to Gαs)^59^. Gαi:β-arrestin comes in close contact to the AP2 adapter complex^24^. Blocking the interaction can impair ERK phosphorylation after GPCR agonist treatment, although interestingly not through all receptors^23,24^. The current evidence suggests that Gαi:β-arrestin interactions can facilitate important cellular signaling events, but that other GPCR signaling pathways independent of this Gαi:β-arrestin interaction are clearly critical for many GPCR signaling transduction events, as certain responses are retained in the absence of β-arrestin.

Taken together, our findings provide an important advance in understanding the mechanistic interplay between G protein and β-arrestin transduction pathways. Further experiments are needed to define how this direct association between two critical mediators of GPCR signaling regulates specific cellular pathways and cellular functions.

## Acknowledgements

We thank Robert J. Lefkowitz, Tom Pack, Meredith Skiba, and Dylan Eiger for thoughtful feedback throughout this work; Kevin Zheng, Nour Nazo, Elizabeth Wren, and Anthony Banks for laboratory assistance. We thank Timothy Adekoya and Ricardo Richardson from North Carolina Central University for the AGS3 constructs. We thank Jonathan Javitch for the D2R-ARB construct. We thank Marc Caron for the MOR construct. We thank Ann Decker from RTI International for providing WW36 and WW38 reagents. This work was supported by 5F31HL152656-03 (C.Y.L.), 1R01GM122798 (S.R.), Mandel Scholar Award (S.R.), the CVRC CARE fellowship (A.G.), the American Heart Association Predoctoral Fellowship 23PRE1019796 (U.P.), The Dermatology Foundation (J.S.S.), K08AR084617 (J.S.S.), the Fujifilm Corporation (A.N.D.), the Smith Family Foundation (A.C.K.), and R01CA260415 (A.C.K.). Authors declare no competing interests. All data are available in the manuscript or the supplementary materials. Data and materials will be fully available upon reasonable request. Constructs designed in this study and that originated in the Rajagopal Lab (table S1) are available from S.R. under a materials transfer agreement with Duke University. Constructs designed in this study and that originated in the Kruse Lab (table S1) are available from A.C.K. under a materials transfer agreement with Harvard University. Graphical figures were created in part using BioRender.com.

## Author contributions

Conceptualization, C.L, J.S.S, S.R; Methodology, C.L, J.S.S, S.R; Investigation, C.L, J.S.S, T.K, A.G, E.M.M, A.S.H, I.C, U.P, F.K, G.G.T, S.R; Writing – Original Draft, C.L, J.S.S; Writing – Reviewing & Editing, C.L, J.S.S, T.K, A.G, I.C, E.M.M, U.P, A.C.K, S.R; Supervision and Funding Acquisition, J.S.S, A.C.K, S.R.

## Materials and Methods

### DNA Plasmid generation

Constructs were generated via conventional cloning methods including restriction digest cloning and overlap cloning. G protein and β-arrestin mutants were generated using Quikchange Lightning mutagenesis (Agilent). Gαi1Nanoluc was synthesized by IDT gBlock (Integrated DNA Technologies, Inc). All constructs that were created, purchased, and received as gifts are denoted in acknowledgements.

### Bacterial strains

XL-1 Blue ultracompetent E. coli, (Agilent), XL-10 Gold ultracompetent E. coli (Agilent) and NEB 5-alpha competent E. coli (New England Biolabs) were used to express and clone constructs.

### Cell culture and transfection

HEK293T cells were maintained in minimum essential medium containing 10% fetal bovine serum and 1% penicillin-streptomycin. Cells were stored in humidified incubator and grown at 37 °C. Cells were transiently transfected by an optimized calcium phosphate60, Fugene, or polyethylenimine (PEI) method61 as previously described. For the calcium phosphate transfection, cell culture media was replaced 30 minutes prior to transfection. DNA constructs were diluted into 90 µL of sterile water. Afterwards, 10 µL of 2.5 M calcium chloride was added to the diluted DNA and mixed. Next, 100 µL of 2xHEPES-buffered saline solution (10 mM D-Glucose, 40 mM HEPES, 10 mM potassium chloride, 270 mM sodium chloride, 1.5 mM disodium hydrogen phosphate dihydrate) was added to the solution dropwise, incubated for 30 seconds-1.5 minutes, and then added to the cells. For Fugene or PEI transfections, a 3:1 ratio of DNA to Fugene or PEI (w/w) was utilized. Briefly, DNA constructs were diluted to 100 µL in Opti-MEM (GIBCO). The appropriate amount of PEI was diluted to 100 µL in Opti-MEM and added to the DNA Opti-MEM. The DNA:PEI solution was mixed, incubated at room temperature for 15-20 minutes, and then added to cells. All transfections were carried out in 6-well cell culture dishes (Costar) unless otherwise denoted.

### NanoBiT Assays and FKBP-FRB Heterodimerization

Experiments were performed as previously described^23^. HEK293T cells were transiently transfected using calcium phosphate. For Gαi:β-arrestin complex formation assays, 500 ng SmBiT-β-arrestin2, 100-200ng of Gα-LgBiT, and 1000-2000 ng of untagged receptor were used. For recruitment assays with SmBiT-β-arrestin2 or Gα-LgBiT, 500-1000 ng of LgBiT or SmBiT tagged receptor was utilized respectively. 18-24 hours post transfection cells were swapped into a starvation medium (clear minimal essential media (MEM) supplemented with 1% penicillin and streptomycin, 1% anti-mycotic anti-biotic, 1% FBS, 10mM HEPES, 1% sodium pyruvate) and seeded into a white, clear bottom, 96-well plate at 50,000-100,000 cells/well. After overnight incubation, starvation medium was removed and replaced with 80 μL of assay buffer containing 20 mM HEPES and 3 μM of coelenterazine-h diluted into Hanks Balanced Salt Solution buffer without calcium or magnesium (HBSS). After 15 minutes, cells were analyzed for basal luminescence at 485 nm for 3 reads at 0.2 msec per well in a BioTek Synergy-Neo2 plate reader set to 37 °C. Following baseline reads, cells were stimulated with either 20 μL of 5X ligand diluted in the assay buffer, or 20 μL of vehicle (assay buffer) and then read for an additional 30 reads. For NanoBiT assays, the net change in luminescence was analyzed as a percent increase over vehicle treatment and normalized to basal luminescence reads. Some assays were additionally normalized to max signal of indicated treatment group.

For NanoBiT FKBP-FRB heterodimerization assays, cells were transfected via 3:1 PEI transfection with 1000 ng Lyn-FRB, 500 ng of SmBiT-β-arrestin2-FKBP and 100-200 ng Gα-LgBiT. 18-24 hours post transfection cells were swapped into a starvation medium and seeded into a white, clear bottom, 96-well plate at 50,000-100,000 cells/well. After overnight incubation, starvation medium was removed and replaced with 80 μL of assay buffer containing 20 mM HEPES and 3 μM of coelenterazine-h diluted into Hanks Balanced Salt Solution buffer without calcium or magnesium (HBSS). After 15 minutes, cells were analyzed for basal luminescence at 485 nm for 3 reads at 0.2 msec per well in a BioTek Synergy-Neo2 plate reader set to 37°C. Following baseline reads, cells were stimulated with either 20 μL of 5X 2.5 µM Rapamycin stock diluted in the assay buffer as previously described29, or 20 μL of vehicle (assay buffer) and then read for an additional 30 reads. The net change in luminescence was analyzed as a percent increase over vehicle treatment and normalized to basal luminescence reads. Plates were read at 37°C with the standard Rluc 485nm emissions filter with 3 baseline reads followed by 30 post-stimulation reads. Rapamycin treated cells were given 500 nM Rapamycin final concentration.

### BRET Assays & NanoBiT BRET Assays

Experiments were performed as previously described^23^. Briefly, constructs were transiently transfected using calcium phosphate or PEI into HEK 293T cells in a 6 well plate. For NanoBRET, 50 ng of Gα-Nanoluc, 500 ng β-arrestin-mKO, and 2000 ng of untagged receptor was used. For internalization assays, optimized amounts of DNA for MyrPalm-mVenus and receptor tagged with RlucII were used (amounts ranged from 500-1000 ng receptor-RlucII and 1000-1500 ng of MyrPalm-mVenus). For β-arrestin recruitment assays, β-arrestin-mKO and receptor-RlucII were used (amounts ranged from 500-1000 ng recep-tor-RlucII and 1000-1500ng of β-arrestin-mKO). For TRUPATH assays62, 1000 ng of untagged receptor, Gαi1-Rluc8, Gβ3, and Gγ9-GFP2 was used. For AGS3 NanoBRET assays, 50 ng of Gα-Nanoluc, 500 ng β-arrestin-mKO, 2000 ng of untagged receptor, and 2000 ng AGS3 were used. For NanoBiT BRET, 100ng of Gαi-LgBiT, 500ng of Gβ1, 500ng Gγ2. 500ng, SmBiT-β-arrestin2, and 2000ng V2R-mKO. After incubating overnight at 37°C, cells were seeded into Costar 96-well plates at 50,000-100,000cells/well in starvation media (see NanoBiT Assay for details). After overnight incubation, starvation medium was removed and replaced with 80 μL of assay buffer containing 20 mM HEPES and 3 μM of coelenterazine-h diluted into HBSS without calcium or magnesium. After 5 minutes, cells were stimulated with appropriate ligands and treatments and analyzed via the following plate reader protocols:

For receptor internalization BRET, β-arrestin recruitment BRET, and NanoBRET assays, plates were read using the BioTek Synergy-Neo2 plate reader set to 37°C with either the standard Rluc 480nm emissions filter and a mKO 542nm long-pass emission filter (Agilent). After stimulation, plates were read 30 times with a 0.5 second integration time. Raw BRET ratio was calculated via dividing the mKO signal by the luminescence signal. ΔNet BRET was calculated by subtracting the average Raw BRET signal of the vehicle wells from the Raw BRET signal of the stimulated wells.

For TRUPATH, plates were read using the BioTek Synergy-Neo2 plate reader or a Glomax reader (Promega) set to 37 °C and a Rluc8 400nm emissions filter and GFP 510nm emissions filter. After stimulation, plates were read 4-6 times. Raw BRET ratio was calculated via dividing the GFP2 signal by the luminescence signal. ΔNet BRET was calculated by subtracting the average Raw BRET signal of the vehicle wells from the Raw BRET signal of the stimulated wells.

For NanoBiT BRET assays, plates were read on a Berthold Mithras LB940 warmed to 37 °C with standard Rluc 485 nm emission filter and a custom mKO 542 nm long-pass emission filter (Chroma Technology Co. Bellows Falls, VT). Plates were read three times to gather basal luminescence and then stimulated with the appropriate concentration of ligand. Afterwards, cells were read for an additional 13 reads to monitor response. ΔNet BRET was calculated by subtracting the average Raw BRET signal of the vehicle wells from the Raw BRET signal of the stimulated wells.

### AlF_4_ Assays

Cells were transfected via calcium phosphate and plated into 96-well plates as previously denoted in the BRET methods. On the day of assay reading, starvation media was replaced with 60 μL of assay buffer containing 20 mM HEPES and 3 μM of coelenterazine-h diluted into HBSS without calcium or magnesium. Three basal reads were taken prior to stimulation on the BioTek SynergyNeo2 plate reader at 37°C using Rluc 480nm emissions filter and a YFP 530nm long-pass emission filter (Agilent). Cells were then treated with 20 µL containing 60 µM of AlF_3_ and 10 mM NaF to generate 60µM of AlF_4_ ^-^ in each well. Cells were incubated for 30 minutes and then read another three times with the reading protocol described above to monitor BRET after treatment. Following basal reads, 20 µL of AVP at various concentrations was added to monitor dose response between 1 pM-10 µM AVP final. After stimulation, the plate was read for an additional 30 reads. AlF_3_ and NaF were purchased from Sigma-Aldrich (St. Louis, MO).

### Recombinant β-arrestin-1 and β-arrestin-2 expression and purification

N-terminally Protein-C tagged rat β-arrestin-1 393X lacking cysteine residues or N-terminally Protein-C tagged bovine β-arrestin-2 392X was purified from BL21 DE3 E. coli. A pMCSG9 expression plasmid encoding maltose binding protein preceded by a hexahistidine tag and followed by a tobacco etch virus cleavage site was appended to the N-terminus of both β-arrestin expression vectors. 10 mL starter cultures were grown in TB supplemented 4% (v/v) glycerol and 100 μg/mL ampicillin to an OD600 of ~2 at 37 °C. Bacteria were cooled to 20 °C for ~2 hours, induced with 200 μM IPTG, and grown overnight at 20 °C. Pellets were harvested and stored at −80 °C until purification. For purification, bacterial pellets were lysed in 50 mM MOPS pH 7.5, 300 mM NaCl, 2 mM DTT lysis buffer with the addition of protease inhibitor tablets and 0.5 mL of 3 mg/mL lysozyme added per liter of culture. After stirring for ~30 minutes at 4 °C, lysates were subsequently passed through a LM10 Microfluidizer Processor three times. Insoluble components were removed via centrifugation (50,000 g for 30 min), and supernatant was syringe filtered through an 0.22 micron filter and passed over Nickel-NTA resin by gravity flow. The column was then washed with 20 column volumes of lysis buffer and eluted with lysis buffer with the addition of 200 mM imidazole. Elution was dialyzed overnight in 50 mM MOPS pH 7.5, 300 mM NaCl, 2 mM DTT with the addition of TEV protease. In the morning, the dialyzed sample was passed over a nickel gravity column twice to remove the cleaved MBP, and then diluted with 50 mM MOPS pH 7.5, 2 mM DTT to a final salt concentration of 100 mM NaCl. Here, purification strategies for β-arrestin-1 and β-arrestin-2 diverged based on a modified protocol from the Gurevich laboratory^63^. β-arrestin-1 was further purified by ion exchange via loading over a HiTrap QXL Column (Cytiva), washed with MOPS pH 7.5, 100 mM NaCl, 100 μM TCEP and eluted with a gradient elution over 40 CV with 50 mM MOPS pH 7.5, 1 M NaCl, and 100 μM TCEP (estimated NaCl concentration at elution peak of 110 mM). β-arrestin-2 was further purified by loading first over a HiTrap QXL column (Cytiva) linked in tandem with a HiTrap SP FF column (Cytiva). After loading, the Q column containing impurities was removed. The SP FF column was then washed with MOPS pH 7.5, 100 mM NaCl, 2 mM DTT and then eluted with a gradient elution over 40 CV with 50 mM MOPS pH 7.5, 1 M NaCl, and 2 mm DTT (estimated NaCl concentration at elution peak of 300 mM). Protein purity was confirmed by SDS–PAGE. Protein was further concentrated, and 10% v/v glycerol was added to the sample. Protein was flash frozen in liquid nitrogen and stored at −80°C until use.

### Recombinant non-lipidated Gαi expression and purification

Human Gαi1 lacking the first three amino acids to remove the lipidation residue were cloned into a modified pET51b expression vector. Gαi1 was preceded by a 10x histidine tag and followed by a 3C cleavage site. 10 mL starter cultures were grown in TB supplemented 4% (v/v) glycerol and 50 μg/mL kanamycin to an OD600 of ~2 at 37 °C. Bacteria were cooled to 20 °C for ~2 hours, induced with 200 μM IPTG, and grown overnight at 20 °C. Pellets were harvested and stored at −80°C until purification. For purification, cells were resuspended in lysis buffer (20 mM HEPES pH 7.4, 100 mM NaCl, 10 mM MgCl _2_, 5 mM imidazole, 5 mM β-mercaptoethanol, 20 μM GDP) at a ratio of 3 mL per gram of cell pellet. Protease inhibitor tablets (1 per 150 mL) and benzonase (1 μL per 150 mL) were added. Cells were lysed using an LM10 homogenizer following a 20–30 minute preincubation on ice. Lysates were clarified by centrifugation at 50,000 × g for 30 minutes at 4°C and filtered through glass fiber filters. The clarified lysate was applied to 2 mL of Ni-NTA resin pre-equilibrated with lysis buffer. The column was washed with 10 CVs of Wash Buffer 1 (20 mM HEPES pH 7.4, 250 mM NaCl, 10 mM imidazole, 5 mM βME, 2 mM MgCl_2_, 10 μM GDP), followed by 10 CV of Wash Buffer 2 (same as Wash Buffer 1 with 20 mM imidazole). Protein was eluted using 4–8 CV of Elution Buffer (20 mM HEPES pH 7.4, 100 mM NaCl, 300 mM imidazole, 5 mM βME, 2 mM MgCl_2_, 10 μM GDP). Elution was monitored using Bradford reagent. The eluate was concentrated to ~12 mL using a 30 kDa cutoff centrifugal filter and incubated with 3C protease at a 1:50 (w/w) estimated protein ratio. The sample was dialyzed overnight at 4°C in 2 L of dialysis buffer (20 mM HEPES pH 7.4, 100 mM NaCl, 5 mM βME, 1 mM MgCl_2_, 10 μM GDP).

Following 3C protease cleavage, the sample was flowed over a second Ni-NTA column to remove uncleaved protein and 3C protease. The flow-through, containing cleaved Gα protein, was collected. The column was further washed with dialysis buffer and subsequently eluted to recover any residual bound proteins. Samples were analyzed by SDS-PAGE. Pooled fractions containing Gαi1 were treated with a phosphatase mix for 1 hour on ice: 5 μL λ phosphatase, 2 μL Quick CIP, 1 μL Antarctic phosphatase, and 1 mM MnCl_2_. The sample was diluted 1:1 in Ion Exchange Dilution Buffer (20 mM HEPES pH 7.4, 5 mM βME, 1 mM MgCl_2_, 10 μM GDP) to a final NaCl concentration of 50 mM.

Ion exchange chromatography was performed using a Mono Q 10/100 column (Cytiva). The sample was loaded via sample pump, washed with 5 CV of Buffer A (20 mM HEPES pH 7.4, 50 mM NaCl, 1 mM TCEP, 1 mM MgCl_2_, 10 μM GDP), and eluted with a linear gradient from 0–40% Buffer B (same as A with 1 M NaCl) over 40 CV. Relevant fractions were pooled, analyzed by SDS-PAGE, concentrated, and stored with 10% glycerol at −80°C.

### Pull downs

Fresh anti-Protein C resin was prepared as a 1:1 slurry in buffer (20 mM HEPES pH 7.4, 100 mM NaCl, 1 mM MgCl_2_, 100 μM TCEP, 2 mM CaCl_2_). For each condition, 25 μL of resin (equivalent to 50 μL slurry) was transferred to a 1.5 mL microcentrifuge tube, spun at 3,500 × g for 2 min at 4 °C, and washed with 1 mL of Buffer. Tubes were shaken gently for 10 minutes in the cold room, followed by a second wash and 5-minute shake.

Protein complexes containing Prc-β-arrestin-1-393X and non-lipidated Gαi1 were assembled at a final concentration of 25 μM on ice for 45 minutes, then incubated with either no treatment, 20 μM GDP, 20 μM GTPγS, or 0.25 units of apyrase (New England Biolabs) for 45 minutes, then equilibrated with antiProtein C resin for 30 minutes in the cold room gently shaking. Resin was pelleted at 3,500 × g for 2 min at 4°C and the supernatant was collected. Beads were washed once with buffer, spun again under the same conditions, and residual buffer was carefully removed using a narrow-31-gauge needle. Proteins were eluted in 40 μL of Elution Buffer (20 mM HEPES pH 7.5, 100 mM NaCl, 1 mM MgCl_2_, 100 μM TCEP, 5 mM EDTA, 0.2 mg/mL Protein C peptide) by shaking for 20 minutes in the cold room, followed by centrifugation at 3,500 × g for 3 min at 4°C. The supernatant was collected using a narrow-31 gauge needle. Eluates were mixed with SDS loading buffer and resolved by SDS-PAGE on 10-well gels at 200 V for 35 minutes. Gels were stained with 1x One-Step Blue Protein Gel Stain (Biotium) and imaged on a Bio-Rad Gel Doc XR+ under Coomassie gel image protocol.

### [35S]-GTPγS binding assays

Gα GTPγS binding assays were described previously^64,65^. Briefly, Gαi1 or myristoylated-Gαi2 (100 nM) were incubated with or without a mixture of β-arrestin-1-393X and β-arrestin-2-392X (400 nM each) at 25 oC in GTPγS binding buffer (20 mM HEPES, pH 8, 100 mM NaCl, 2 mM DTT, 1 mM EDTA, 10 mM MgCl2 and 0.05% (m/v) C12E10 and 10 µM [35S]GTPγS (SA 4,000 cpm/pmol). Aliquots were taken from biological triplicate reactions at indicated time points (3, 6, 10, 15, 30, 60, 90 min), quenched in GTP quench buffer (20 mM Tris, pH 7.7, 100 mM NaCl, 10 mM MgCl2, 1 mM GTP, and 0.08% (m/v) C12E10), and filtered onto BA-85 nitrocellulose filters (GE Healthcare Life Science). Filters were washed (20 mM Tris, pH 7.7, 100 mM NaCl, 2 mM MgCl2), dried and subjected to scintillation counting.

### Ligands

AVP, dopamine, AngII, DAMGO, BAM22, Quinpirole, Fentanyl, and isoproterenol, tiotropium were purchased from Sigma-Aldrich (St. Louis, MO). PZM21, SR17018, and TRV130 (Oliceridine) were purchased from MedKoo Biosciences (Cary, NC). TRV120023 was synthesized by Genscript (Piscataway, NJ). WW36, WW38, were provided via a collaboration with RTI International (Research Triangle Park, NC). Stock solutions of AVP, AngII, DAMGO, and BAM22 were prepared according to the manufacturer’s specifications. Isoproterenol, WW36, and WW38 were dissolved in dimethyl sulfoxide (DMSO) and stored in a desiccator cabinet. Fentanyl was provided pre-solubilized in ethanol vials (1mg/mL). All drug dilutions were performed with BRET medium or cell culture medium. PTX was obtained from List Biological Laboratories (Campbell, CA) and stored at 4°C. All compound stocks were stored at −20°C until use.

### Sequence Alignments

Human protein sequence alignments for Gαi1, Gαi2, and Gαi3 were obtained from GproteinDb66 and visualized on SnapGene (San Diego, CA)

### Data Analysis

Data were analyzed via Excel (Microsoft, Redmond, WA) and graphed in Prism10.0 (GraphPad, San Diego, CA). Log agonist versus stimulus with three parameters (span, baseline, and EC50) was used to determine dose-response curves for applicable data sets. Comparisons of ligand or mutant constructs in concentration and time response assays were conducted via a two-way analysis of variance (ANOVA) unless otherwise noted. Most experiments were conducted three or more times. Some replicates represented positive or negative controls. Statistical tests were two-sided and Bonferroni analysis was corrected for multiple comparisons unless otherwise noted. Lines represent mean values and error bars represent SEM unless otherwise noted.

