## Supplemental Figures for "β-arrestin recruitment facilitates a direct association with G proteins"

### Slide 1
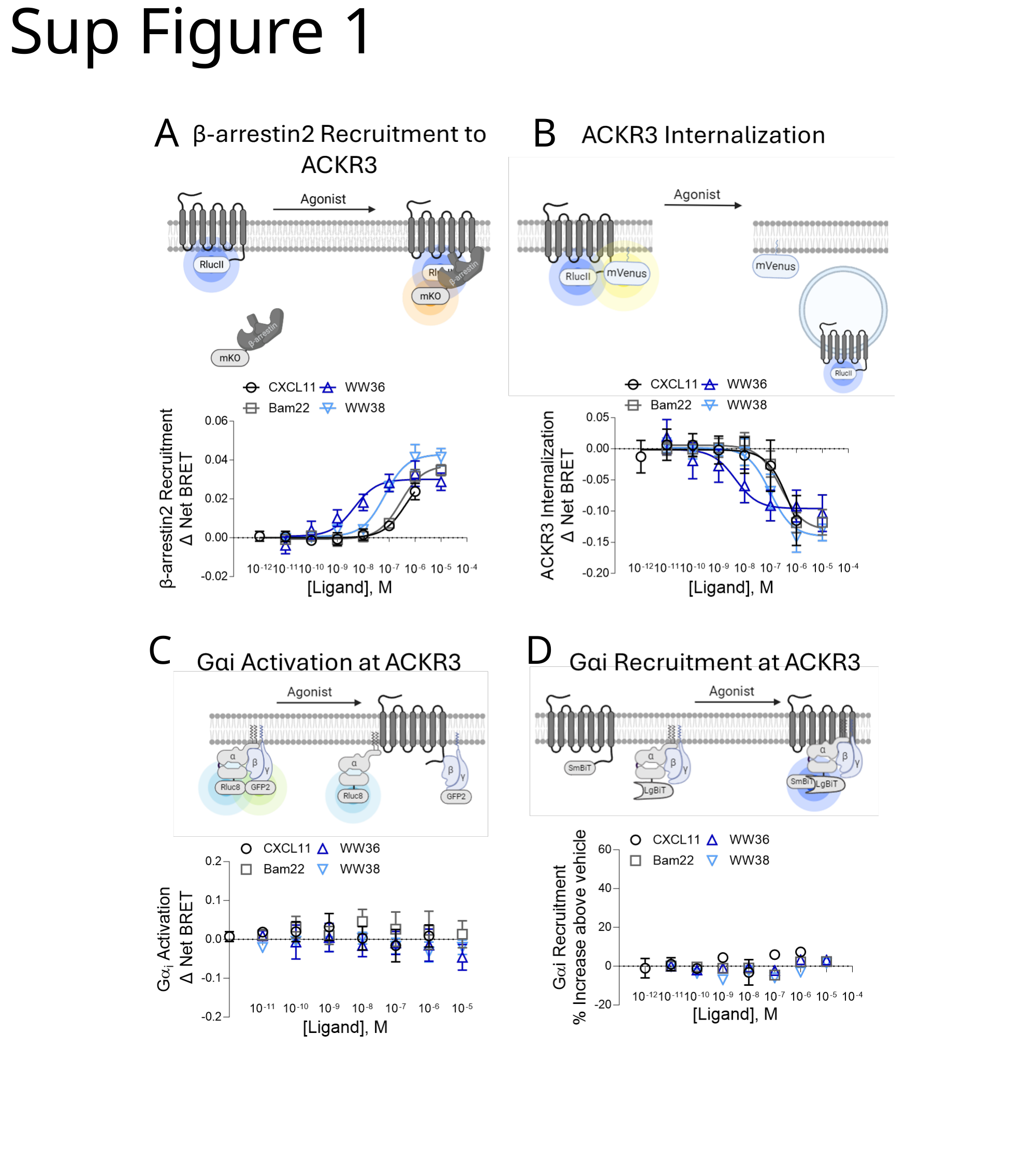

#### Sup Figure 1
A
B
C
D

### Slide 2
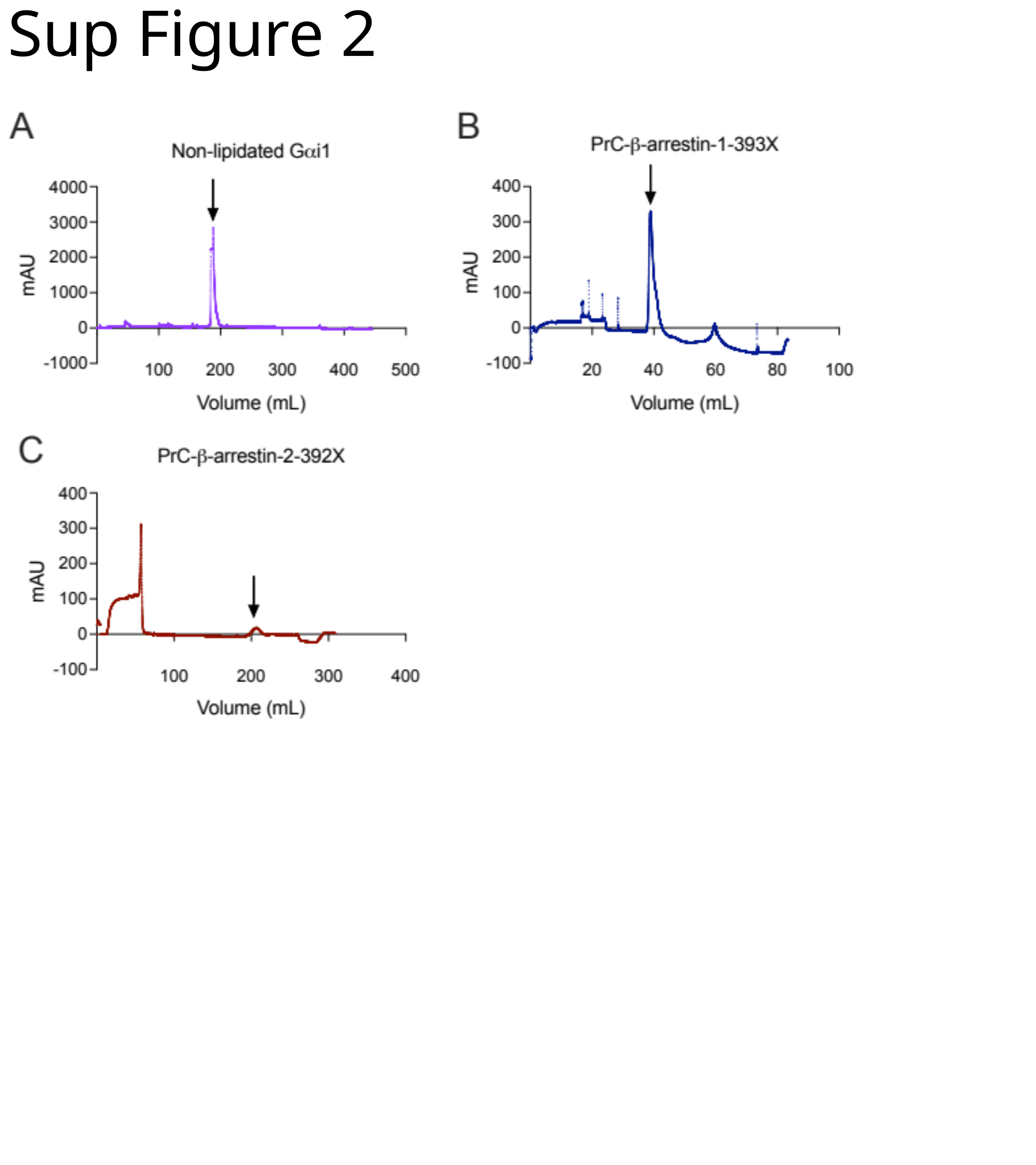

Sup Figure 2

### Slide 3
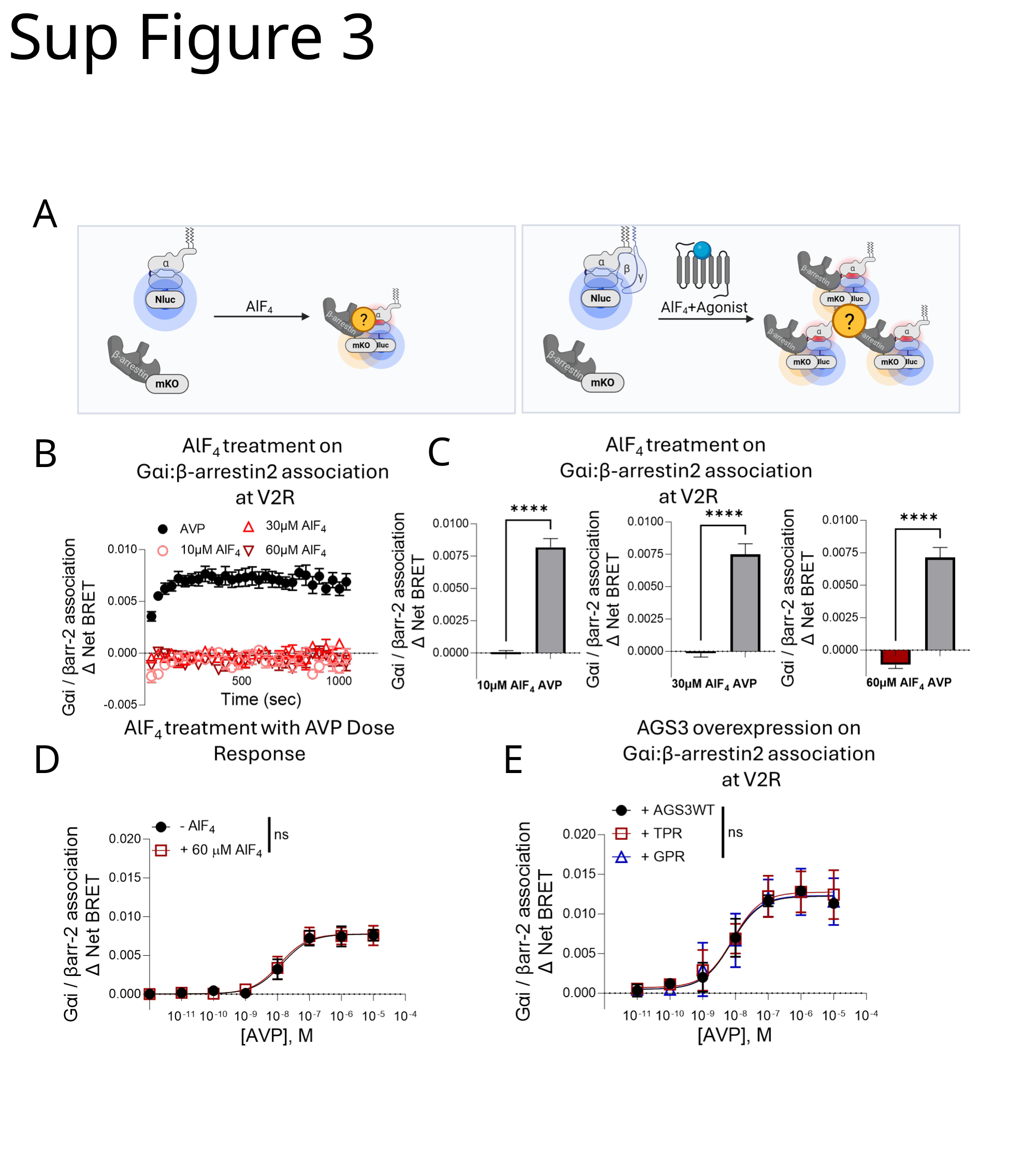

#### Sup Figure 3
A
C
B
E
D

### Slide 4
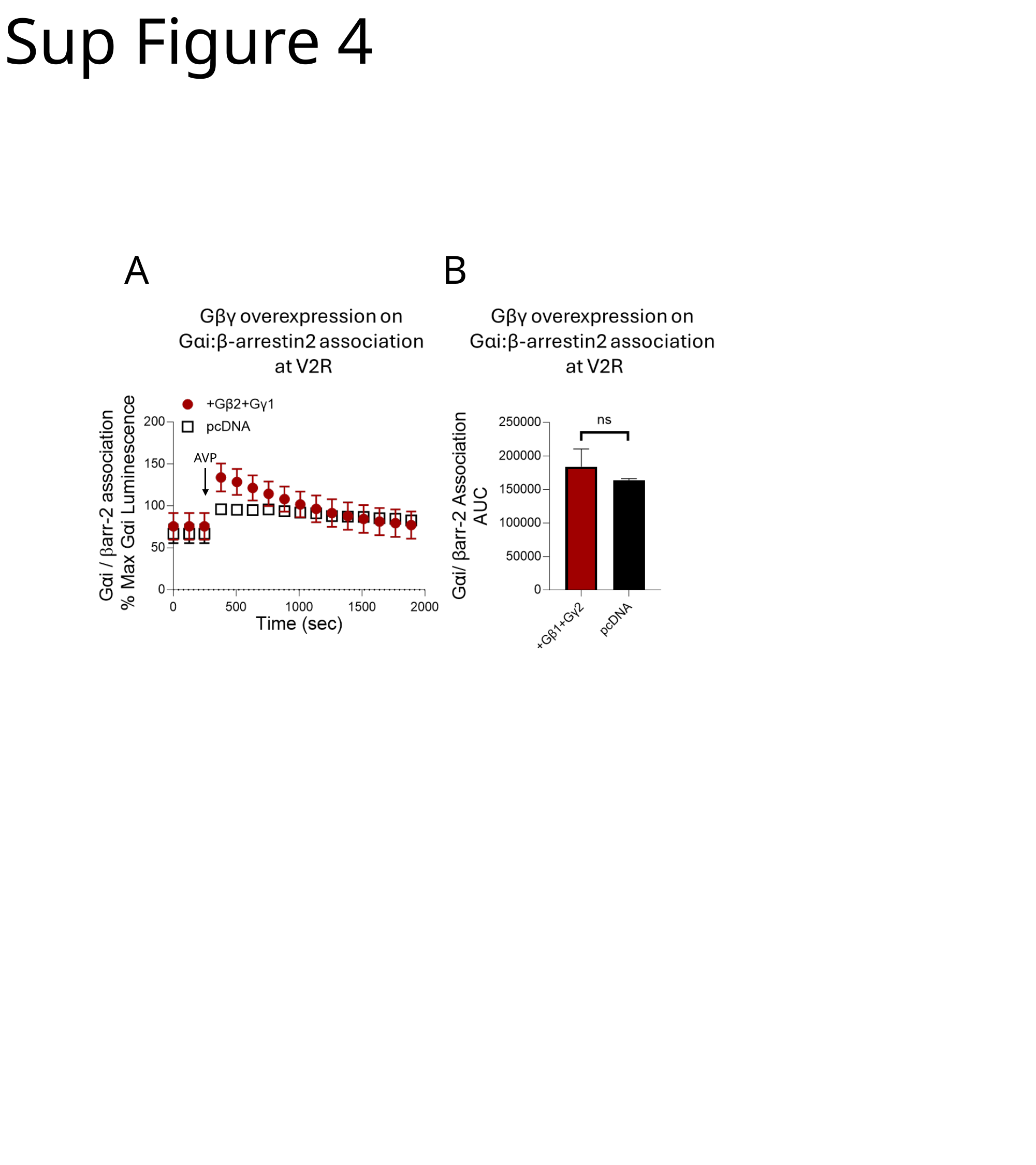

#### Sup Figure 4
B
A
AVP

### Slide 5
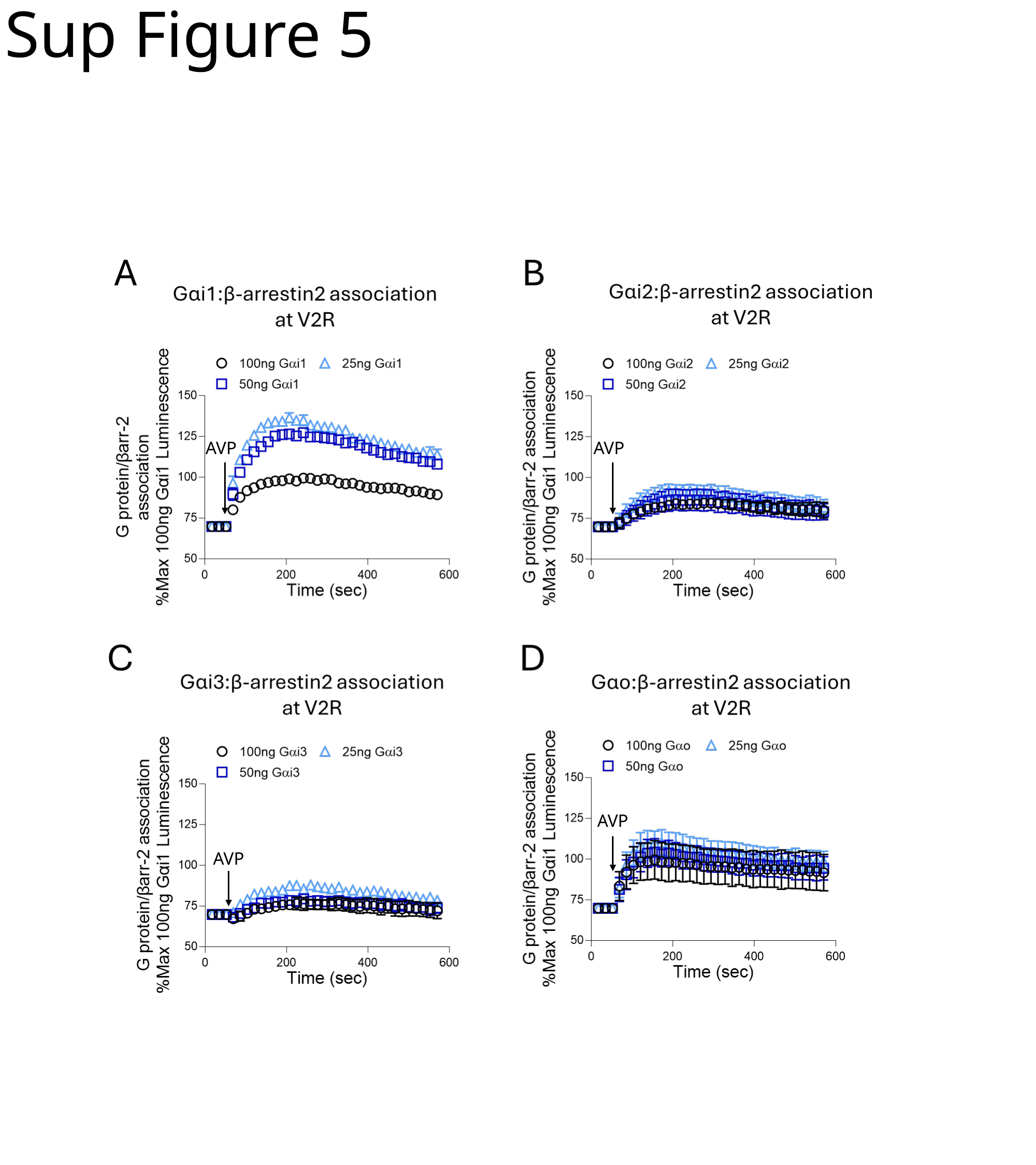

#### Sup Figure 5
AVP
AVP
AVP
AVP

### Slide 6
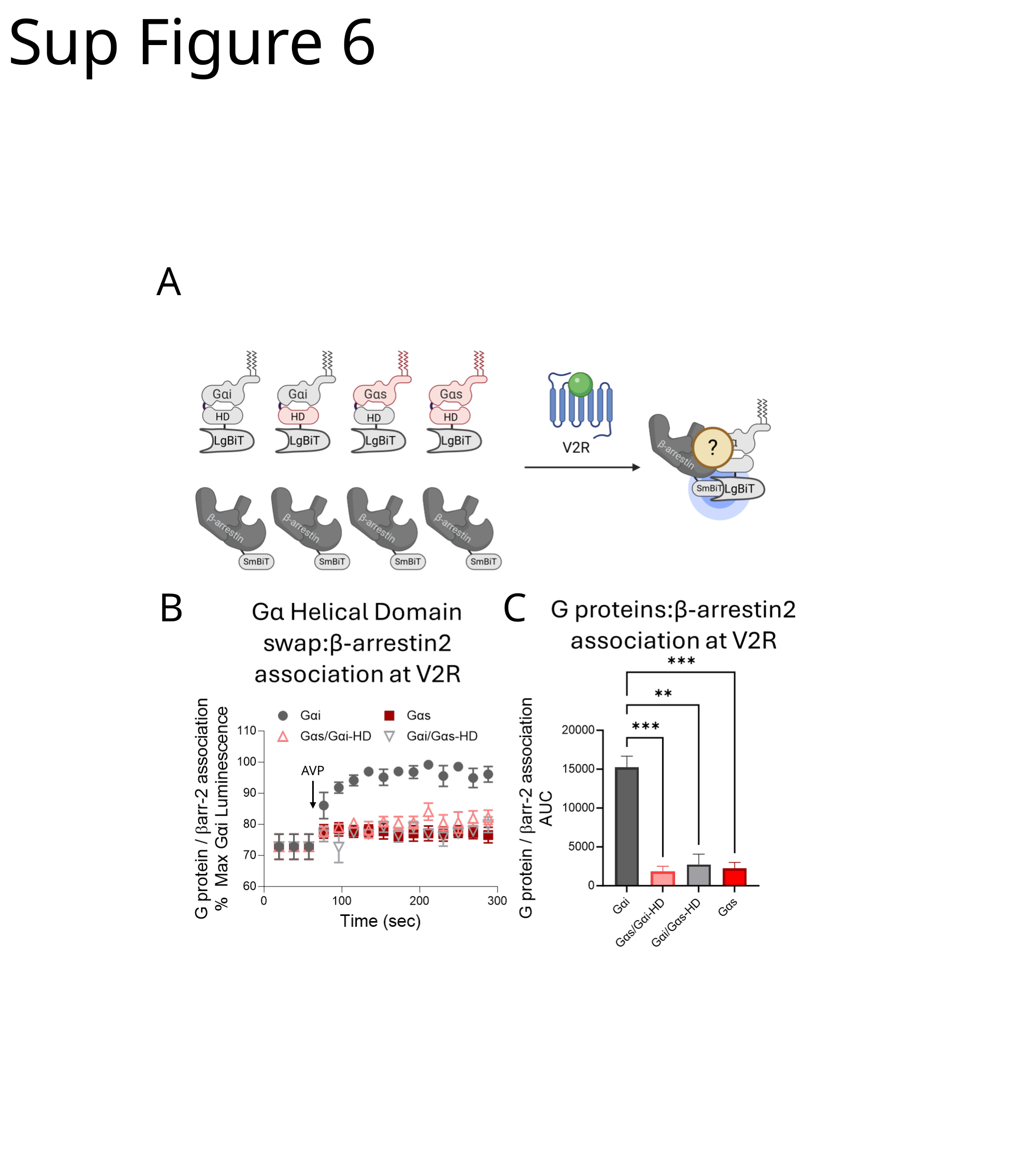

#### Sup Figure 6
A
B
C
AVP
